# Topographic connectivity reveals task-dependent retinotopic processing throughout the human brain

**DOI:** 10.1101/2020.07.30.228403

**Authors:** Tomas Knapen

## Abstract

The human visual system is organized as a hierarchy of maps that share the topography of the retina. These retinotopic maps have been identified throughout the brain, but how much of the brain is visually organized remains unknown. Here we demonstrate widespread stable visual organization beyond the traditional visual system. We analyzed detailed topographic connectivity with primary visual cortex during moviewatching, rest, and retinotopic mapping experiments to reveal that visual-spatial representations are warped by experimental condition and cognitive state. Specifically, traditionally visual regions alternate with default mode network and hippocampus in preferentially representing the center of the visual field. This visual role of hippocampus would allow it to implement sensory predictions by interfacing between abstract memories and concrete perceptions. These results indicate that pervasive sensory coding facilitates the communication between far-flung brain regions.

---

Our entire experience of the world is ultimately based on impressions arriving through the senses. In our dominant sensory modality, vision, processing is *retinotopic*: organized according to the layout of the retina^1^. That is, neigbouring locations in the brain represent neighbouring locations in the visual field. Retinotopic mapping experiments leverage sparse visual stimulation during fixation, allowing researchers to relate the elicited brain responses to visual space and delineate retinotopic maps in the brain^2,3^. Yet, in everyday life visual inputs are not sparse, and naturalistic vision is characterized by continuous eye movements and dynamic cognitive demands. It is therefore likely that charting especially high-level visual function falls outside the scope of traditional retinotopic mapping experiments.

Retinotopic processing throughout the brain can be identified based on topographically specific connectivity with V1^4–6^, the first visual region of the cerebral cortex (Fig 1a). There are several distinct advantages to assessing visual processing by means of retinotopic connectivity (RC), in which we explain responses throughout the brain in terms of the spatial pattern of activation on the surface of V1. First, RC is robust in the face of eye movements, because its reference frame is fixed in the brain and not the outside world. Second, because V1 harbors a map of visual space, RC patterns throughout the brain can be translated back into visual space. In effect, RC allows us to project the retinotopy of V1 into the rest of the brain. Lastly, since brain responses are explained as a function of ongoing activations, RC can be estimated for any experimental paradigm. Thus, RC can be used to compare detailed visual-spatial processing across experiments and cognitive states.

**Figure 1.**
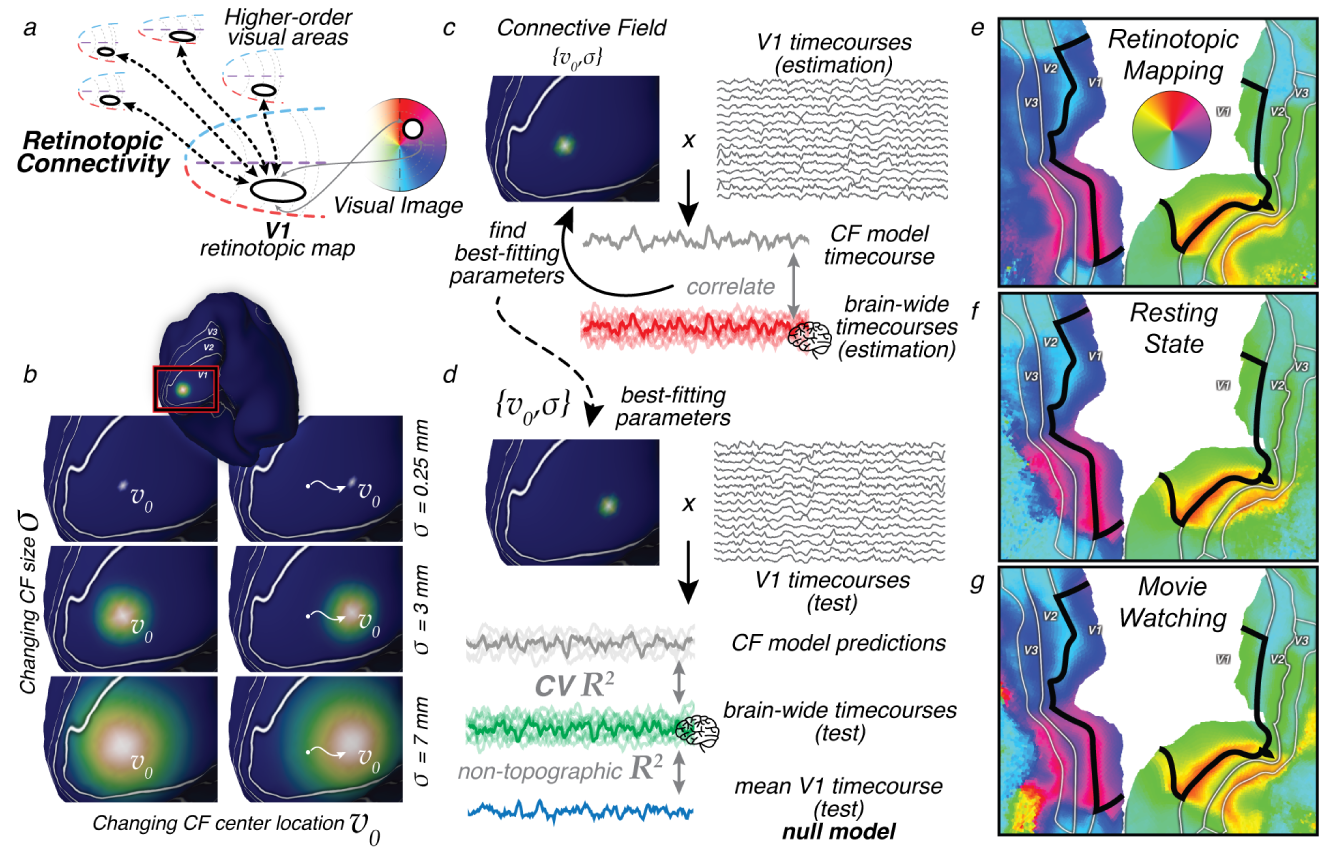
**a**. Visual processing in higher-order brain regions reveals itself through spatially specific retinotopic connectivity with V1. V1’s map of visual space allows us to translate retinotopic connectivity to representations of visual space. **b**. Retinotopic connectivity is quantified by modeling responses throughout the brain as emanating from Gaussian Connective Fields on the surface of V1. Example Gaussian CF model profiles with different size (*σ*) and location (*v*_0_) parameters are shown on the inflated surface. **c**. Predictions are generated by weighting the ongoing BOLD signals in V1 with these CF kernels. We find the optimal CF parameters to explain signal variance for all locations throughout the brain (non-V1 timecourses). **d**. Cross-validation procedure. We assess CV prediction performance of the CF model a left-out on test data set, and correct for the performance of a non-topographic null model, the average V1 timecourse. **e,f,g**. CF modeling results can be used to reconstruct the retinotopic structure of visual cortex. Polar angle preferences outside the black outline are reconstructed solely based on their topographic connectivity with V1, within the black outline. CF modeling was performed on data from retinotopic mapping, resting state and movie-watching experiments separately. Visual field preferences are stable across cognitive states, as evidenced by the robust locations of polar angle reversals at the borders between V2 and V3 field maps. Additional retinotopic structure visualizations in Supplementary Figure 1 & 2.

We performed RC analysis on the Human Connectome Project (HCP) 7T dataset of 174 subjects in which data were collected during retinotopic mapping, resting state, and movie watching experiments. This allowed us to identify previously unknown visual-spatial processing throughout the brain, quantify how visual space is flexibly represented - even in brain regions not traditionally considered visual - and reveal how these visual representations depend on cognitive state.

A parsimonious computational model for RC posits that responses arise from a localized Gaussian patch on the surface of V1 (see Figure 1b), its connective field^5^(CF). We fit the CF model by comparing CF model time-course predictions to ongoing BOLD response time-courses throughout the brain (see Figure 1c&d). Translating the best-fitting CF parameters into visual field locations reveals the structure of visual field maps in V2, V3, and beyond (Figure 1e). Retinotopic maps resulting from retinotopic mapping, movie-watching, and resting state acquisitions are similar, with their borders in the same location. This means that in low-level visual cortex the structure and strength of this retinotopic connectivity is both stable across participants and robust against variations in experimental task, stimulation and cognitive state.

Does this RC extend beyond the lower levels of the visual system, and if so, how does it depend on the different cognitive states evoked in different experiments? Indeed, Figure 2a shows that significant portions of movie-watching BOLD fluctuations throughout the cerebral cortex are explained as resulting from RC. That is, more than half of the cerebral cortex, including large swaths of the temporal and frontal lobes, shows significant topographically-specific connectivity with V1. Interestingly, the local strength of RC depends heavily on the experiment (Fig 2b). During movie watching, mainly ventral visual and temporal brain regions exhibit RC. This retinotopic connectivity likely reflects object identity-related processing and multisensory integration^7^. Conversely, during resting-state scans the default-mode network (DMN) shows stronger RC, which may reflect endogenous mental imagery during mind-wandering^8^.

**Figure 2.**
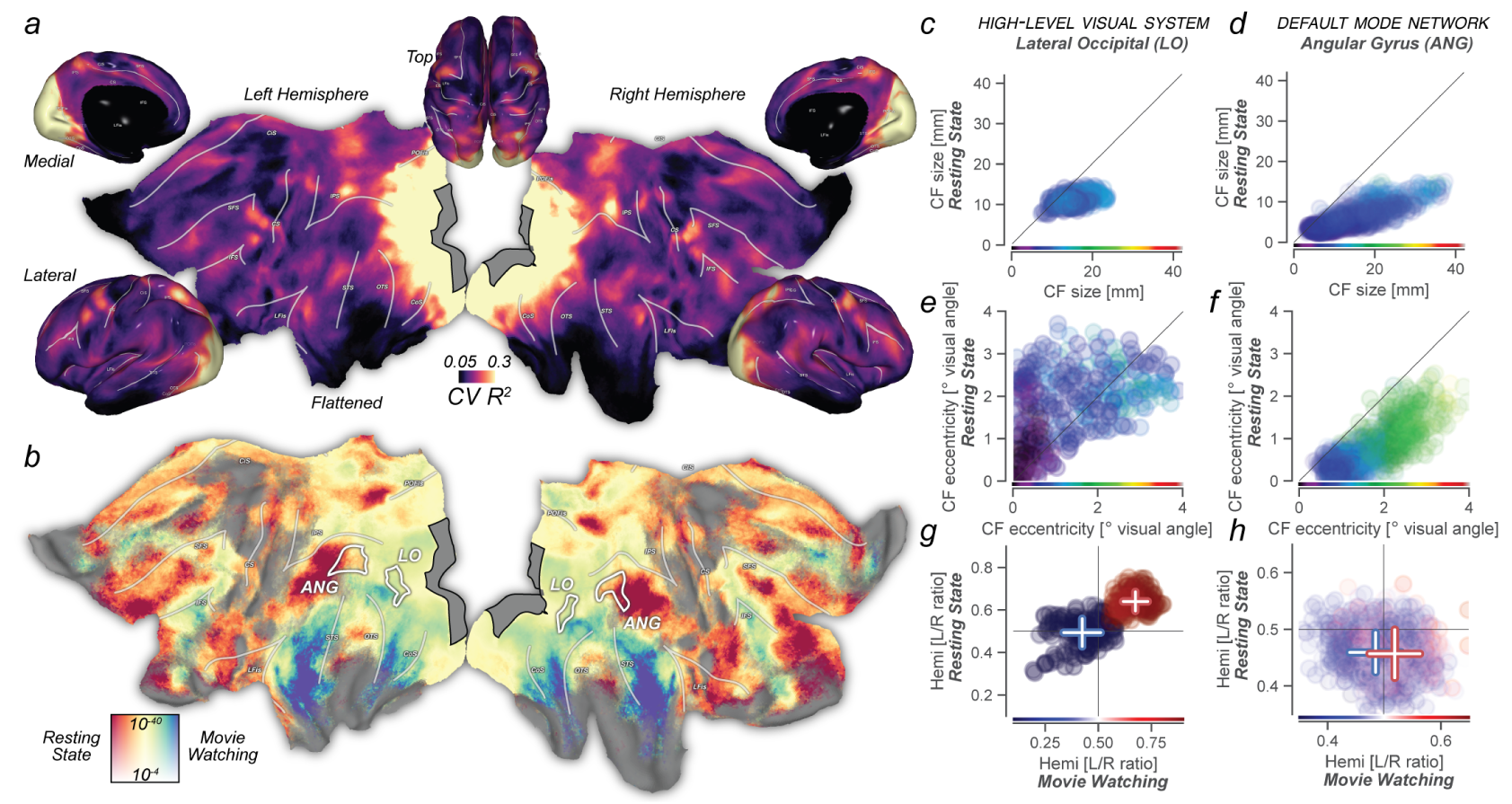
**a**. Retinotopic connectivity during movie watching explains significant amounts of variance throughout the brain. The gray region contains all possible CF center locations; all vertex locations that are not black have significant best-fitting CV R2. **b**. Retinotopic connectivity is modulated by cognitive state. The local strength of topographic connectivity to V1 depends on whether participants were engaged in movie watching or endogenous thought. Ventral and lateral visual regions sensitive to object identity and audiovisual processing are more strongly connected to V1 during movie-watching. Topographic connectivity between Default-Mode Network regions and V1 is strongest during resting state. Colormap for resting state vs movie watching represents normalized variance explained ratio (see methods) and ranges from 0.1 to 0.9. Vertical color scale axis represents p-values. **c**. Scatter plots show CF and CF-derived visual field parameters for three experimental conditions. Horizontal and vertical axes represent movie watching and resting state results, respectively. Point color represents parameter values from the retinotopy experiment and is defined on the same range as the scatter plots. Point opacity linearly relates to null-model-corrected R^2^ averaged across experimental conditions. In the high-level visual system, CF-derived eccentricity correlates strongly between conditions. We see marked foveal biases in visual field preference for movie watching and retinotopic mapping experiments relative to resting state. **d**. CF size is fixed and similar during resting state and retinotopic mapping experiments, with CF size variation during movie watching showing a strong correlation in CF size between conditions. **e**. The contralaterality of visual location preference (quantified as average CF hemisphere of origin) are strongly correlated across experimental conditions. **f,g,h**. Like in LO, CF-derived retinotopic representations in the DMN are stable, as evident from high correlations in eccentricity and size between conditions. During resting state, CF become more foveal - a pattern opposite to the cognition-driven shifts of CF spatial positions in LO. During visual stimulation, the DMN represents visual space similarly to high-level visual regions^17^. Supplementary Figures 2 & 3 visualize CF tuning for additional brain regions and CF parameters across experiments. For detailed statistics see Supplementary Table 2.

If retinotopic visual processing is a stable organizational property, visual field preferences derived from one experiment should predict RC from another. The detailed inspection of CF parameters can provide insights into the visual-spatial processing embodied by RC, but also allows us to understand its modulation by cognition. We compare two regions, lateral-occipital (LO) and angular gyrus (ANG), as exemplars of high-level visual and default-mode network (DMN) areas, respectively. In both regions, spatial sampling extent as quantified by CF size, is stable^9^ and precise^5,10^ during both retinotopy and resting state and becomes larger and more variable during movie watching (Fig 2c&d). Sampling extent is strongly correlated between conditions (all linear correlations > 0.36, all *p* < 10^−13^ (LO), > 0.75, all *p* < 10^−37^ (ANG) - full statistics in Supplementary Table 1). This points to stable sampling of retinotopic space in both high-level visual and DMN regions.

In LO, the preferred eccentricity of cortical locations (Fig 2e) is strongly correlated between experimental conditions (linear correlations > 0.48, all *p* < 10^−19^), confirming its well-known retinotopy^9,11^ (Fig 2c). The detailed differences in spatial representations between conditions, however, show the flexibility of LO’s retinotopy: a very foveal bias during retinotopic mapping^11^ gives way to broader coverage of the visual field during resting state and movie watching. The across-condition robustness of ANG eccentricity (Fig 2f) is greater than that of LO (all linear correlations > 0.75, all *p* < 10^−40^). However, ANG shows an opposite pattern of retinotopic flexibility, with visual field preferences being more foveal specifically during the resting state. These foveopetal and foveofugal changes of eccentricity in both LO and ANG mimic the effects of top-down attention^12,13^ and imagination^14^ on visual field representations. This pattern is also similar to what happens to visual items of interest as they are foveated through eye movements^15^, and follows the topographic distribution of feedback related to visual object information^16^ in the absence of eye movements.

V1 represents contralateral visual field locations, allowing us to use the hemisphere of the best-fitting CF as a proxy for visual field representation laterality. In LO, visual representations are strongly contralateral, and strongly correlated between experimental conditions (t-test between hemispheres: all *T*(> 123) > 10, *p* < 10^−^19). In ANG, there is a significant contralaterality bias of visual field representations during retinotopic mapping and movie watching (all *T*(199) > 6, *p* < 10^−^9), but, presumably due to the strong foveal bias of visual field representations, not during resting state (*T*(199) = 0.14, *p* = 0.9). These findings confirm previous reports that the DMN represents visual space similarly to high-level visual brain regions in situations of visual stimulation^17^.

Although it is not generally implicated in traditional vision science experiments, tracer-based connectivity studies place the hippocampal formation at the top of the visual processing hierarchy^18^. Hip-pocampus is thought to implement the interaction between memory-related and sensory processing or imagery^19–22^, leading us to reason that both narrative understanding of naturalistic inputs and internally generated thought should evoke strong RC in hippocampus, over and above its recently discovered contralateral visual field preference during retinotopic mapping^23^ (Supplementary Fig 4).

Applying CF modeling to hippocampus BOLD timecourses (Fig 3a) reveals significant hippocampal-V1 RC in all experimental conditions. We observe a gradient of movie watching vs resting state RC preference along both the medial/lateral and the long, anterior/posterior axes of the hippocampus. This RC gradient corresponds in detail to previously found microstructural and memory-related functional connectivity gradients^24,25^. The high power, quality, and spatial resolution of both functional and anatomical HCP images allow us to quantify cognitive state-dependent RC per hippocampal subfield. We predict, based on earlier findings in both mouse^22^ and humans^26^, that the CA region and uncus of the hippocampus should subserve its connectivity with visual cortex during visual stimulation.

**Figure 3.**
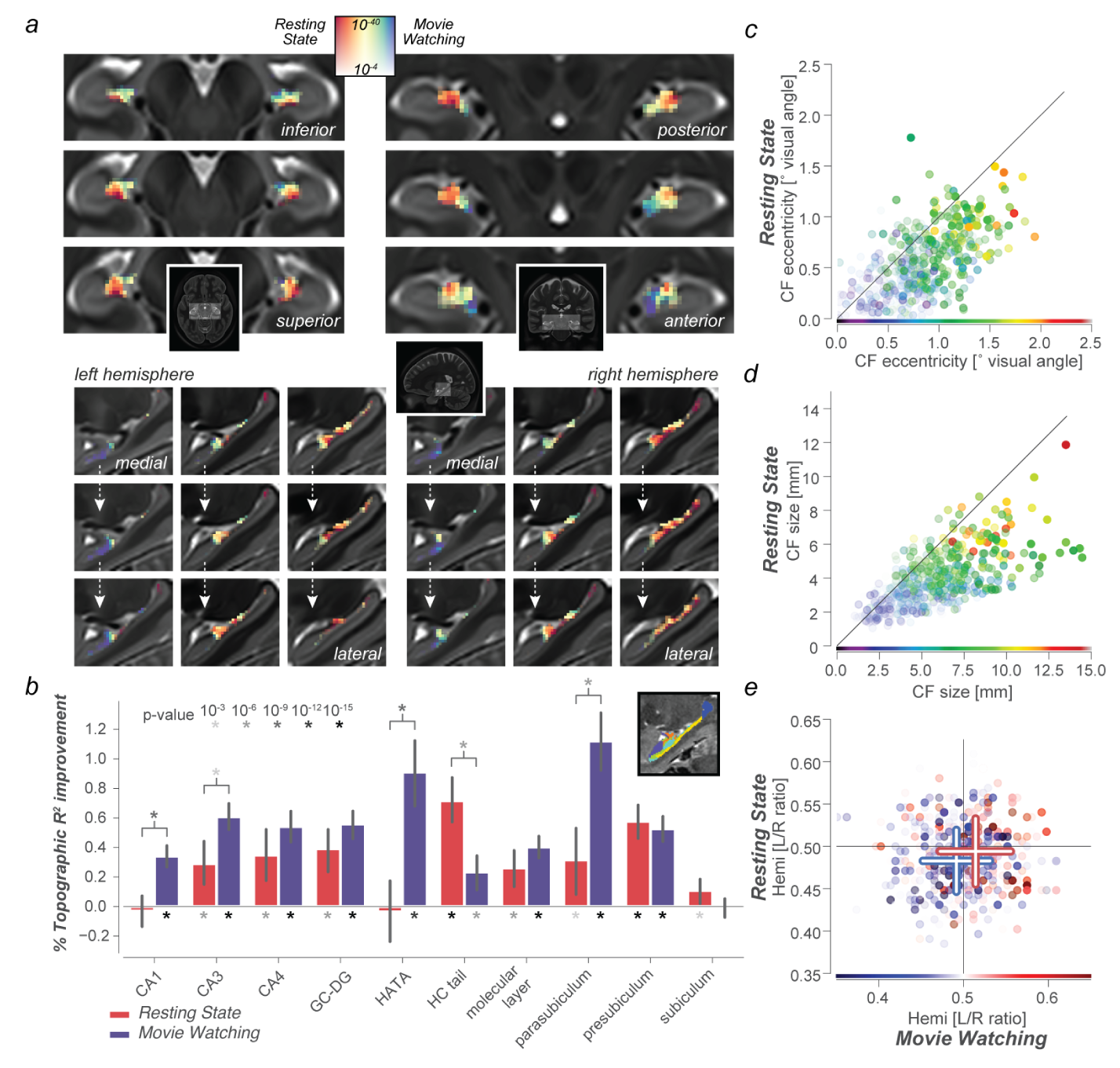
RC is pervasive throughout the hippocampal formation. **a**. Like the cerebral cortex, the hippocampus also displays a gradient of endogenous/exogenous processing, with stronger topographic connectivity in rostral/medial/inferior regions during movie watching, and in caudal/lateral/superior regions during resting state. Colormap for resting state vs movie watching represents normalized variance explained ratio ranges from 0.35 to 0.65. Full lightbox visualization of RC connectivity gradient and visual field parameters in Supplementary Figure 4. **b**. Strength and stability of RC for different cognitive states, for separate hippocampal subfields. Inset shows a sagittal section of a hippocampal subfield segmentation for a single subject. Full statistics given in Supplementary Tables. Eccentricity **c**. and Size **d**. of hippocampal CFs is stable between different cognitive states, with strong correlations between all three conditions (linear CF eccentricity and size correlations > 0.55, *p* < 10^−26^). **e**. Moreover, visual field representations are biased to predominantly represent the contralateral visual hemifield in movie watching, resting state, and retinotopy (t-test between hemispheres, *T* = −4.3, *p* < 2 · 10^−5^, *df* = 173, *T* = −3.4, *p* < 10^−3^, *df* = 173, and *T* = −10.3, *p* < 10^−20^, *df* = 173, respectively). Detailed visualization of visual field representations in hippocampal and thalamic subregions in Supplementary Figure 5.

The strength and cognitive state-dependence of RC varies strongly across hippocampal subfields (Fig 3b). Specifically, RC driven by movie watching is strongest in parasubiculum, CA1, CA3, Dentate Gyrus, and the hippocampal-amygdalar transition area which principally represents the hippocampal uncus. Conversely, resting-state driven RC is strongest in the presubiculum and the hippocampal tail. Focusing on the visual space representations of hippocampus, as quantified by CF model parameters, we see strong correlations between experimental conditions (Fig 3c,d&d). Specifically, the detailed patterns of differences in visual field representations between experimental conditions closely resemble those that occur in the DMN (Fig 2 and Supplementary Fig 3 & 5).

How do these findings relate to other connectivity-based findings? Variations in RC amplitude between experimental conditions are compellingly similar to large-scale connectivity gradients found during resting state^27^. These large-scale gradients result from the push-pull between sensory and attentional brain systems on the one hand, and the DMN on the other^28^. Such countervailing activations and deactivations are thought to mediate a dynamic balance between outward-oriented, stimulus-driven processing on the one hand, and memory-related, endogenously generated processing on the other^29,30^. Importantly, the present work identifies a consistent mode of organization across both memory-related and stimulus-directed processing: in both cognitive states, connectivity is retinotopically organized. Furthermore, we show that the push-pull between DMN and sensory regions in terms of signal amplitude^28^ also involves a trade-off in foveally-biased processing between regions. On the basis of our findings, we propose that the detailed structure of concurrent retinotopically organized activations in visual system, DMN and hippocampus gives rise to interactions between attentional and mnemonic processing^31^.

The present finding that the hippocampus shares a retinotopic mode of organization with much of the rest of the brain, is in line with canonical tracer-based network findings^18^. As hippocampus also entertains world-centric coding of space^32^, this solidifies the notion^19^ that hippocampus is a nexus for the conjunctive coding of both world-centric and sensory reference frames^20,33^.

The stability of RC structure across experiments points to the brain’s use of sensory topography as a fundamental organizing principle to facilitate neural communication between distant brain regions. By casting ongoing brain responses into a common reference frame, RC can facilitate our understanding of neural processing as resulting from canonical computational mechanisms such as divisive normalization^34^. Future work will be able to leverage RC to investigate the computational hierarchy that generates increasingly world-centric representations of space^35^ and culminates in the medial temporal lobe^32,36^.

## Methods

### Human Connectome Project data

HCP 7T fMRI data were used, in conjunction with 3T anatomical MRI images^37^. In total, 2.5 hours from 174 subjects with full data of all 7T experiments were used, sampled at 1.6 mm isotropic resolution and a rate of 1Hz^38^. For all functional analyses, we used the Fix ICA-denoised timecourse data, sampled to the 59k vertex-per-hemisphere `MSMAll` surface and 1.6mm MNI volume formats. These data are freely available from the HCP project website.

### Analysis

Hippocampal subfield segmentation was performed using FreeSurfer, after which the individual subfields were warped to the functional data’s MNI space using the existing HCP warpfields with nearest-neighbour interpolation. High-resolution subfield segmentations were smoothed with a Gaussian of 0.8 mm *σ* to ensure representation of all subfields when resampled to the 1.6mm resolution of the functional images. Anatomical ROI definitions were taken from the multimodal parcellation atlas^39^ for surface data, and FSL’s Jülich histological atlas^40^ for hippocampal ROIs in MNI volumetric space.

All in-house analyses were implemented in python, using scientific python packages. The full list of dependencies for running the analyses, and the code itself, are available on GitHub. The notebooks in this repository allow users to recreate all data visualizations presented here.

Functional data of all three experiments (movie (approx. 1 hour), retinotopy^41^ (30 mins), and resting state (1 hour)) were preprocessed identically, by means of high-pass filtering (3rd order Savizky-Golay filter, 210s period) and z-scoring over time.

To create a template of retinotopic spatial selectivity, we averaged the timecourses of the retinotopic mapping experiment (bar and wedge conditions) across participants, and estimated linear Gaussian population receptive field (pRF) model parameters from

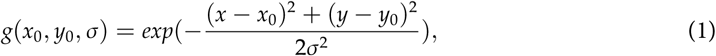

where *x*_0_, *y*_0_ and *σ* are the parameters that define the location (in the cartesian {*x, y*} plane) and size of the pRF, respectively. The fitting procedure consisted of an initial grid fit stage, followed by an iterative fitting stage using the L-BFGS-B algorithm as implemented in `scipy.optimize`. The identical fitting procedure was performed on the hippocampus to create figures of visual-spatial representations shown in Supplementary Figure 4.

Gaussian connective field profiles on the surface are defined for each vertex *v* on the cortical manifold as

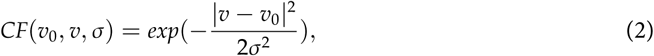

where *v*_0_ is the center location of the connective field, and *σ* is the Gaussian spread in mm on the cortical surface. Diffusion of heat along the surface mesh was simulated and then used to infer geodesic distances |*v* − *v*_0_| between all V1 vertices, as implemented in pycortex^42^ (which was also used for all surface-based visualization). As the two hemispheres are two separate surfaces, this distance matrix was calculated for each hemisphere separately. Only V1 vertices with a within-set R^2^ of >0.2, a peak pRF position inside the stimulus aperture used in the retinotopy experiment, and a positive pRF amplitude in the above pRF analysis served as the center of a candidate CF. These conservative selection criteria were chosen to ensure that CFs are centered on visually responsive vertices within V1, and improves the interpretability of the relation between CF parameters and visual field coordinates. We verified that using the full V1 map as possible CF center yielded similar RC results on the whole. We used a fixed set of candidate sizes, ranging from very small (biased to the center vertex only) to evenly spanning almost the entirety of V1: [0.5, 1, 2, 3, 4, 5, 7, 10, 15, 20, 30, 40, 80] mm *σ* for the Gaussian CF. When releasing this constraint of known topographic organization in the source region, CF modelling and RC estimation exemplify a broader category of analysis techniques in which decomposition of signals based on their local structure represents a very efficient manner of mapping all-to-all correlations between brain activations into a subspace relevant to local neural processing^43^.

The predicted timecourse for each CF was generated by taking the dot product between the CF’s vertex profile and the vertex by timecourses matrix. These CF timecourses were z-scored and correlated with the timecourses throughout the brain, without convolution with a haemodynamic response function. For each location in the brain, we selected the connective field resulting in the highest correlation. Subsequently, we used this specific connective field to generate model timecourses for out-of-set prediction of left-out data. We used a 4-fold scheme for cross-validation. As both the resting state and movie watching consist of 4 separate runs of approximately 15 minutes each, this scheme was implemented to be identical to a leave-one-run-out cross-validation pattern. CF parameters and explained variance measures were then averaged across runs for further analysis.

The resulting proportion of out-of-set explained variance was referenced against the proportion explained variance of a non-spatial null model, in which the timecourses throughout the brain were predicted by the average V1 timecourse. This correction means that although corrected R^2^ values are no longer interpretable as proportion explained variance, they can serve to conservatively assess the presence of true topographic connectivity. Moreover, comparisons between conditions based on these corrected *R*^2^ values explicitly discount changes in non-topographic correlations between brain regions. One-sample t-tests were used to compare corrected out-of-set prediction performance against 0, reported p-values are two-sided. Correlations across vertices, as reported in Supplementary Table 1, were calculated weighted by this corrected R^2^. To quantify a location’s RC preference during resting state vs movie watching, we created a normalized ratio of the *null-model* corrected CV R^2^ values for the respective experimental conditions: 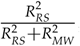, where RS and MW stand for resting state and movie watching, respectively. This measure takes a value between 0 and 1, where 0.5 signifies equal RC in both experimental conditions.

## Acknowledgments

TK was supported by a high-performance computing grant (NWO ENW-17683). Data were provided by the Human Connectome Project, WU-Minn Consortium (Principal Investigators: D. Van Essen and K. Uğurbil; 1U54MH091657) funded by the 16 NIH Institutes and Centers that support the NIH Blueprint for Neuroscience Research; and by the McDonnell Center for Systems Neuroscience at Washington University in St. Louis. The data are available for download at www.humanconnectome.org.

I thank Ed Silson, Chris Baker, Peter Zeidman, Nicholas Hedger, & Eli Merriam for providing comments on an earlier version of the manuscript, and Kendrick Kay, Alex Huth, & Ben Hutchison for fruitful discussions.

## Supplementary Materials

### Supplementary Tables

**Table 1:**
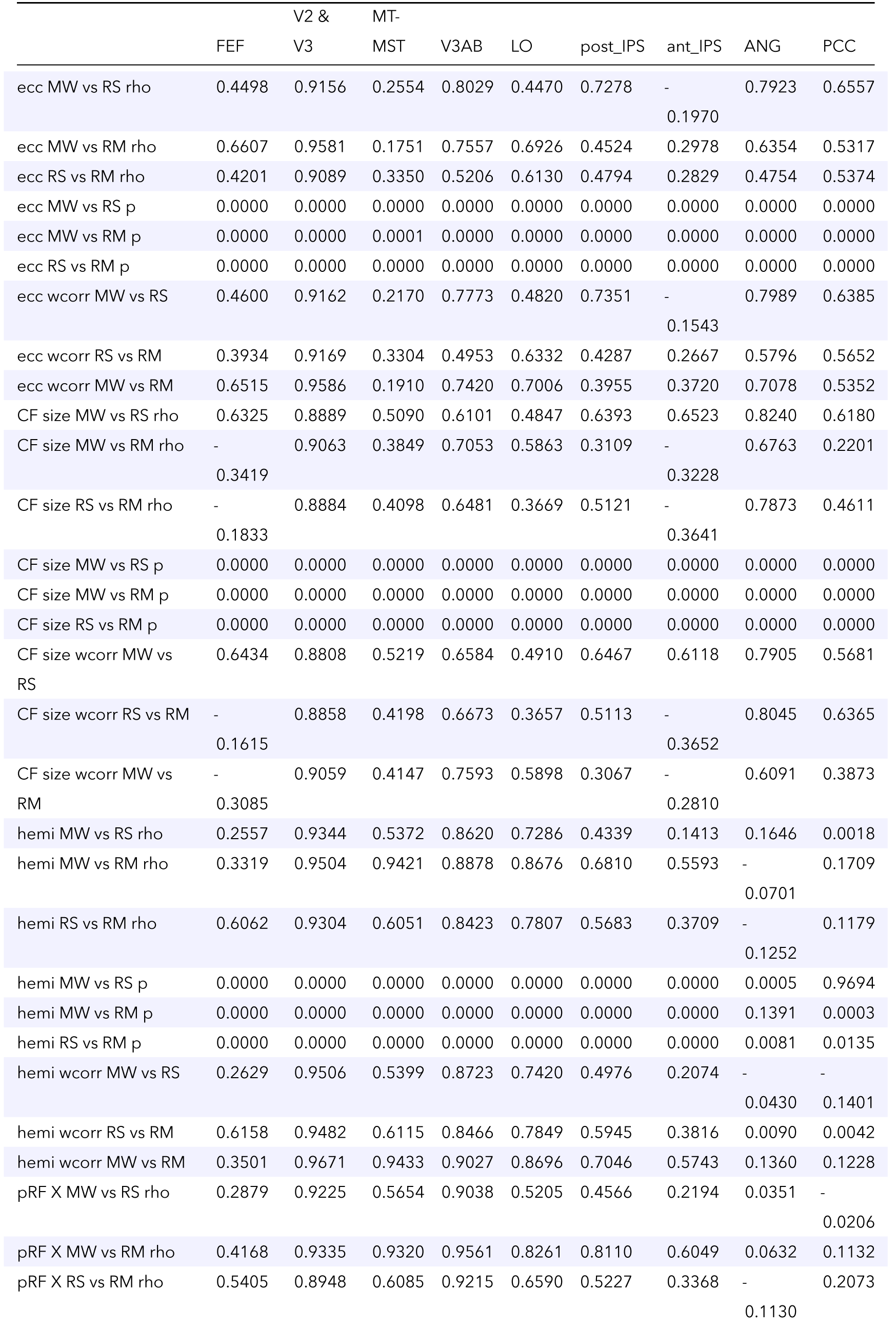
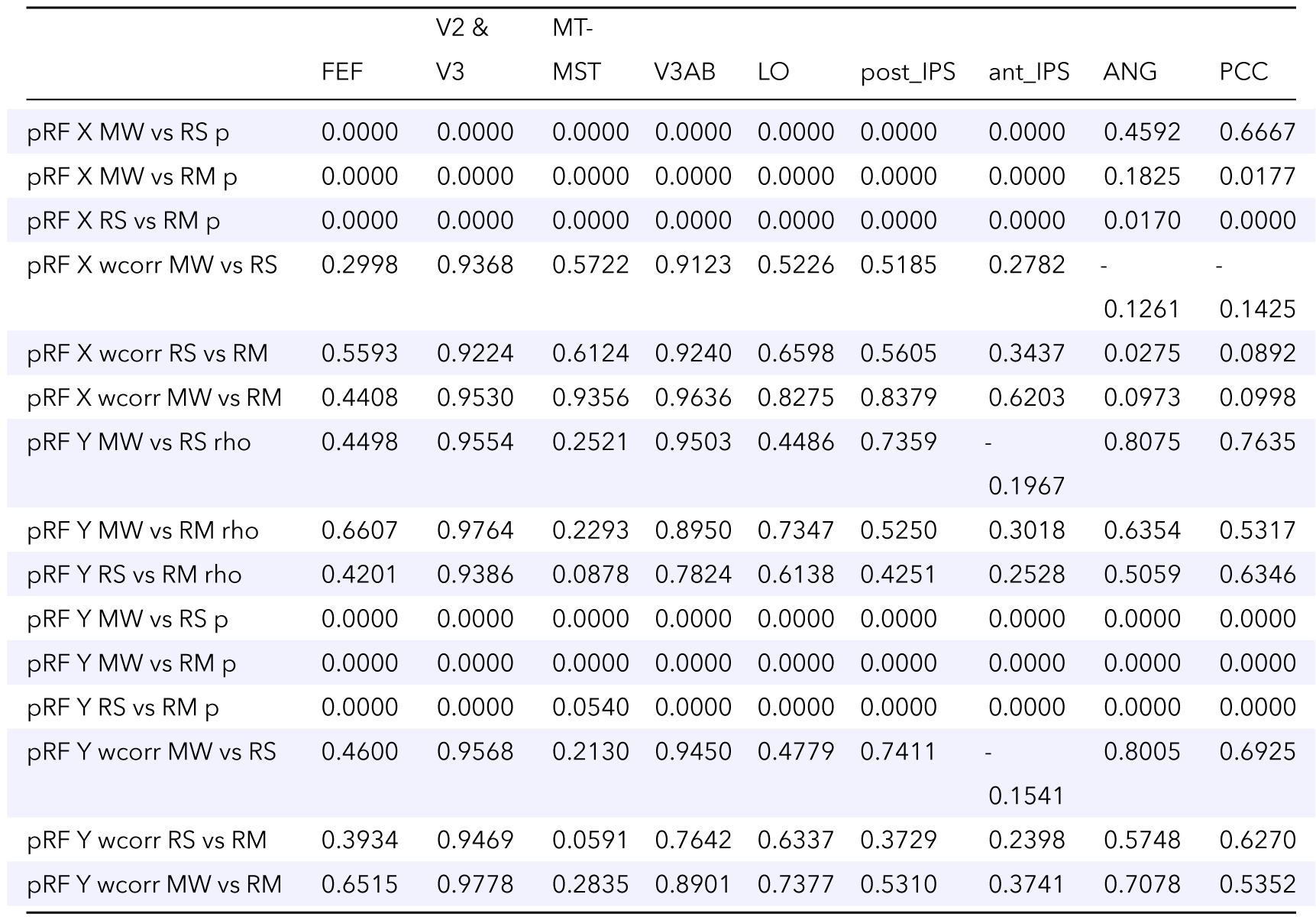
Statistics for CF parameter correlations in cerebral cortex. MW; Movie Watching, RS; Resting State, RM; Retinotopic Mapping. rho; linear correlation coefficient, wcorr; linear correlation coefficient weighted by r-squared.

**Table 2:** Statistics for movie-watching RC in different hippocampal subfields

| Movie watching | T | dof | p-val | cohen-d | BF10 | power |
| --- | --- | --- | --- | --- | --- | --- |
| CA1 | 9.941 | 173 | 5.28584e-19 | 0.754 | 9.507e+15 | 1 |
| CA3 | 13.855 | 173 | 3.77237e-30 | 1.05 | 8.602e+26 | 1 |
| CA4 | 9.897 | 173 | 6.96224e-19 | 0.75 | 7.259e+15 | 1 |
| GC-DG | 11.367 | 173 | 5.0978e-23 | 0.862 | 8.263e+19 | 1 |
| HATA | 7.753 | 173 | 3.67912e-13 | 0.588 | 1.881e+10 | 1 |
| HP_tail | 3.813 | 173 | 9.55131e-05 | 0.289 | 163.061 | 0.984 |
| molecular_layer_HP | 10.848 | 173 | 1.51676e-21 | 0.822 | 2.954e+18 | 1 |
| parasubiculum | 10.813 | 173 | 1.90166e-21 | 0.82 | 2.366e+18 | 1 |
| presubiculum | 11.985 | 173 | 8.74929e-25 | 0.909 | 4.489e+21 | 1 |
| subiculum | -0.35 | 173 | 0.636802 | 0.027 | 0.18 | 0.023 |

**Table 3:** Statistics for resting-state RC in different hippocampal subfields

| Resting State | T | dof | p-val | cohen-d | BF10 | power |
| --- | --- | --- | --- | --- | --- | --- |
| CA1 | -0.574 | 173 | 0.716772 | 0.044 | 0.199 | 0.013 |
| CA3 | 4.016 | 173 | 4.40624e-05 | 0.304 | 335.108 | 0.991 |
| CA4 | 3.888 | 173 | 7.20393e-05 | 0.295 | 211.937 | 0.987 |
| GC-DG | 5.275 | 173 | 1.96592e-07 | 0.4 | 55870 | 1 |
| HATA | -0.344 | 173 | 0.63439 | 0.026 | 0.179 | 0.023 |
| HP_tail | 9.485 | 173 | 9.55299e-18 | 0.719 | 5.593e+14 | 1 |
| molecular_layer_HP | 4.167 | 173 | 2.43291e-05 | 0.316 | 583.993 | 0.994 |
| parasubiculum | 2.695 | 173 | 0.00387086 | 0.204 | 5.605 | 0.851 |
| presubiculum | 9.74 | 173 | 1.89864e-18 | 0.738 | 2.718e+15 | 1 |
| subiculum | 2.585 | 173 | 0.0052737 | 0.196 | 4.267 | 0.824 |

**Table 4:** Statistics for the difference between resting-state and movie watching RC in different hippocampal subfields

|  | T | dof | p-val | cohen-d | BF10 | power |
| --- | --- | --- | --- | --- | --- | --- |
| CA1 | -5.87 | 346 | 1.02077e-08 | 0.629 | 970000 | 1 |
| CA3 | -3.748 | 346 | 0.000208772 | 0.402 | 91.057 | 0.962 |
| CA4 | -1.84 | 346 | 0.0666119 | 0.197 | 0.599 | 0.45 |
| GC-DG | -1.889 | 346 | 0.0597003 | 0.203 | 0.654 | 0.47 |
| HATA | -5.99 | 346 | 5.26794e-09 | 0.642 | 1.818e+06 | 1 |
| HP_tail | 4.955 | 346 | 1.13427e-06 | 0.531 | 11380 | 0.999 |
| molecular_layer_HP | -1.933 | 346 | 0.0541079 | 0.207 | 0.707 | 0.487 |
| parasubiculum | -5.153 | 346 | 4.31793e-07 | 0.552 | 28180 | 0.999 |
| presubiculum | 0.695 | 346 | 0.487632 | 0.074 | 0.149 | 0.107 |
| subiculum | 2.262 | 346 | 0.024307 | 0.243 | 1.366 | 0.616 |

### Supplementary Figures

**Supplementary Figure 1.**
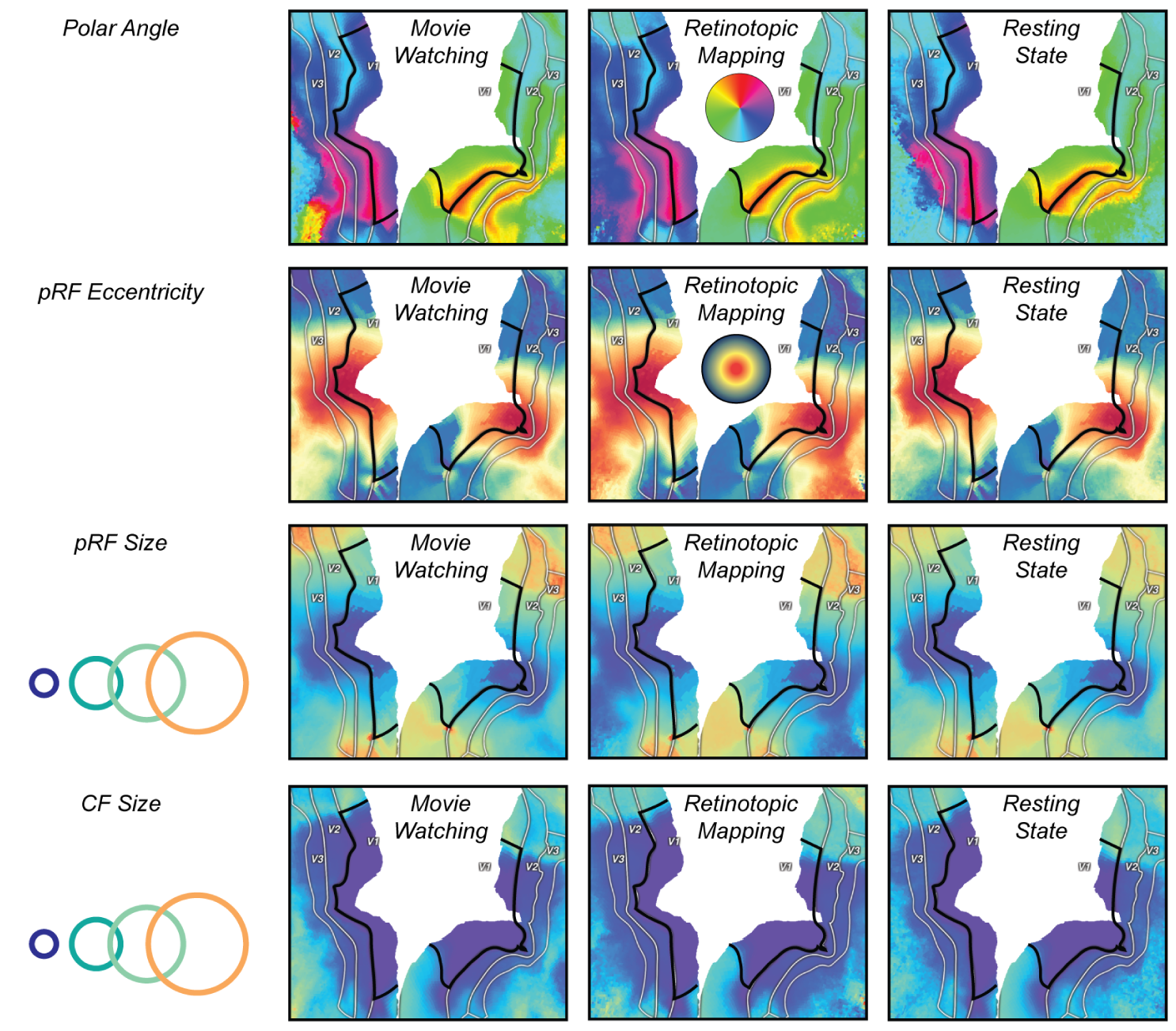
Detailed visualization of retinotopic parameters resulting from CF fit in the visual system. View identical to that of Fig 1 e,f&g.

**Supplementary Figure 2.**
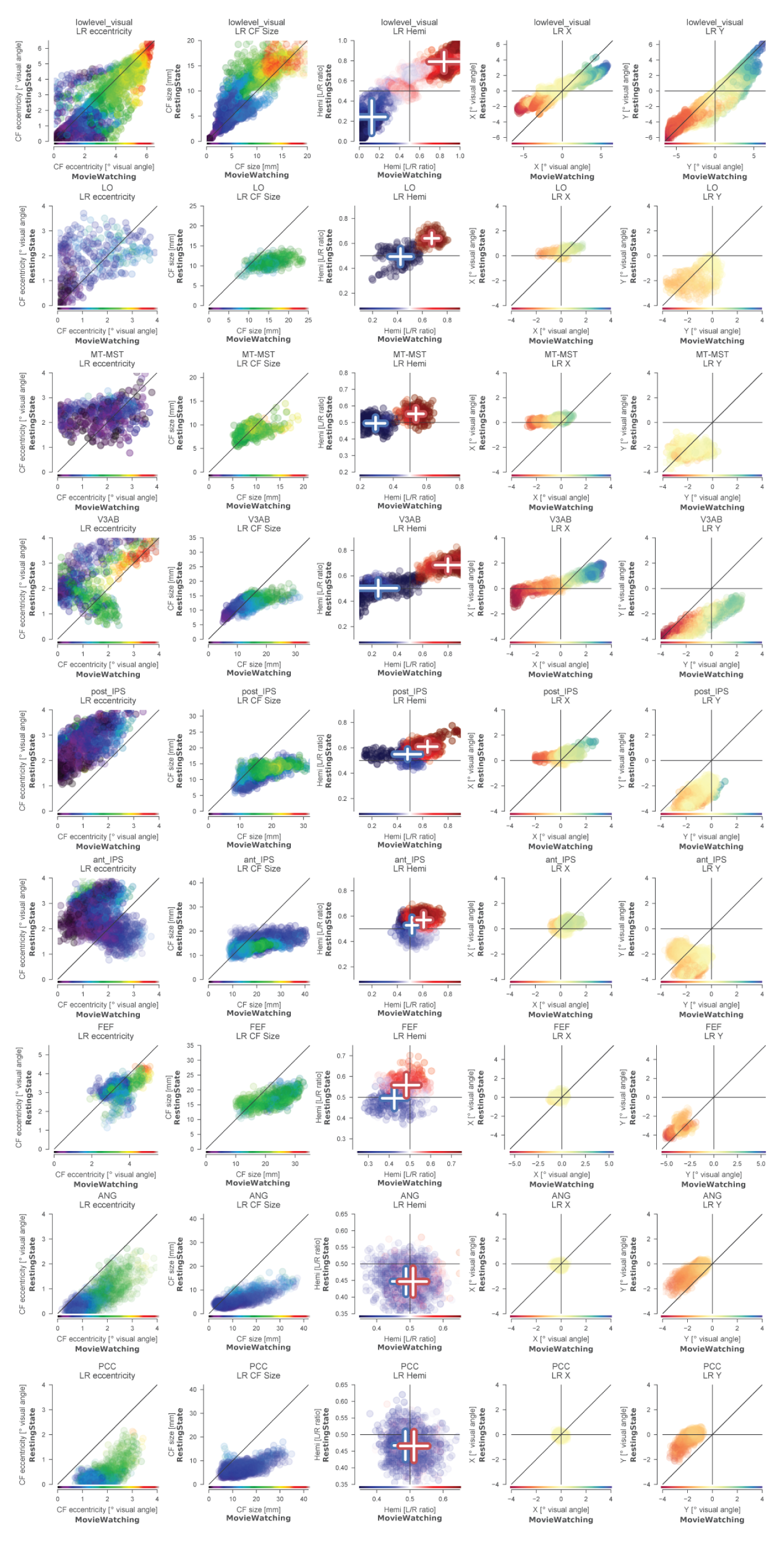
CF parameter correlations between conditions for several visual regions in the dorsal visual processing stream, from lower to higher levels of processing. All ROI definitions are taken from the MMP_HCP atlas^39^. “lowlevel_visual” combines V2 and V3. This figure adds columns for x and y CF/pRF parameters.

**Supplementary Figure 3.**
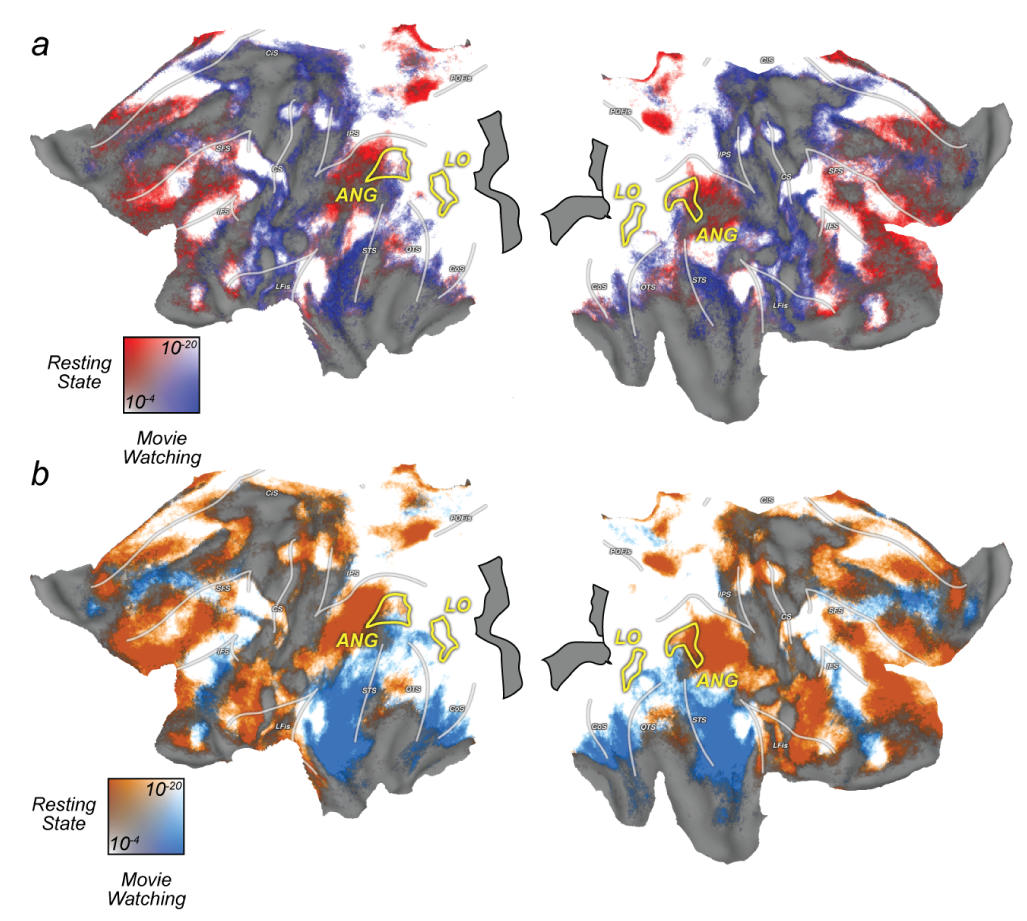
CF fits are stable across experiments. **a**. It is possible to fit the CF model on one experiment, and then predict the BOLD timecourses of a second experiment. Colours represent negative log_1_0 p-values for training on Resting State and testing on Movie Watching (Blue) and viceversa (Red). In most regions (White) prediction across experiments is highly significant in both directions. Reference against the same visualization of within-experiment prediction (**b**.) reveals that the preference of a given location depends on the experiment on which it was tested. This indicates that 1. CF estimates are stable across conditions, and 2. differences in RC between experiments are driven by the strength at which RC is driven in the test, and not the train condition.

**Supplementary Figure 4.**
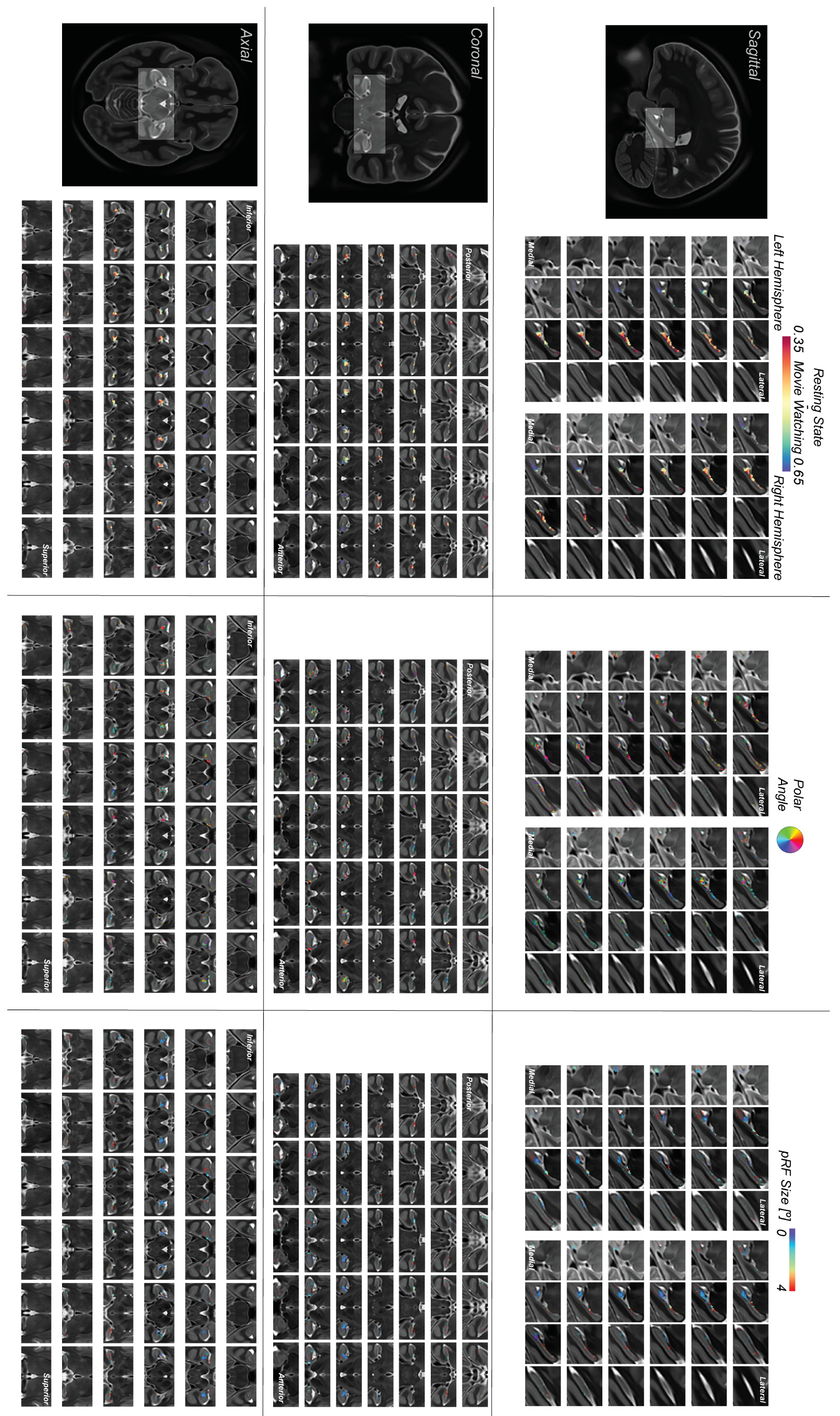
Full zoomed lightbox views of relative strength of hippocampal retinotopic connectivity in Resting State and Movie Watching (left), pRF polar angle (middle) and pRF size (right), were derived from the retinotopic mapping experiment, analyzed identically to V1 (See Fig 1 & methods).

**Supplementary Figure 5.**
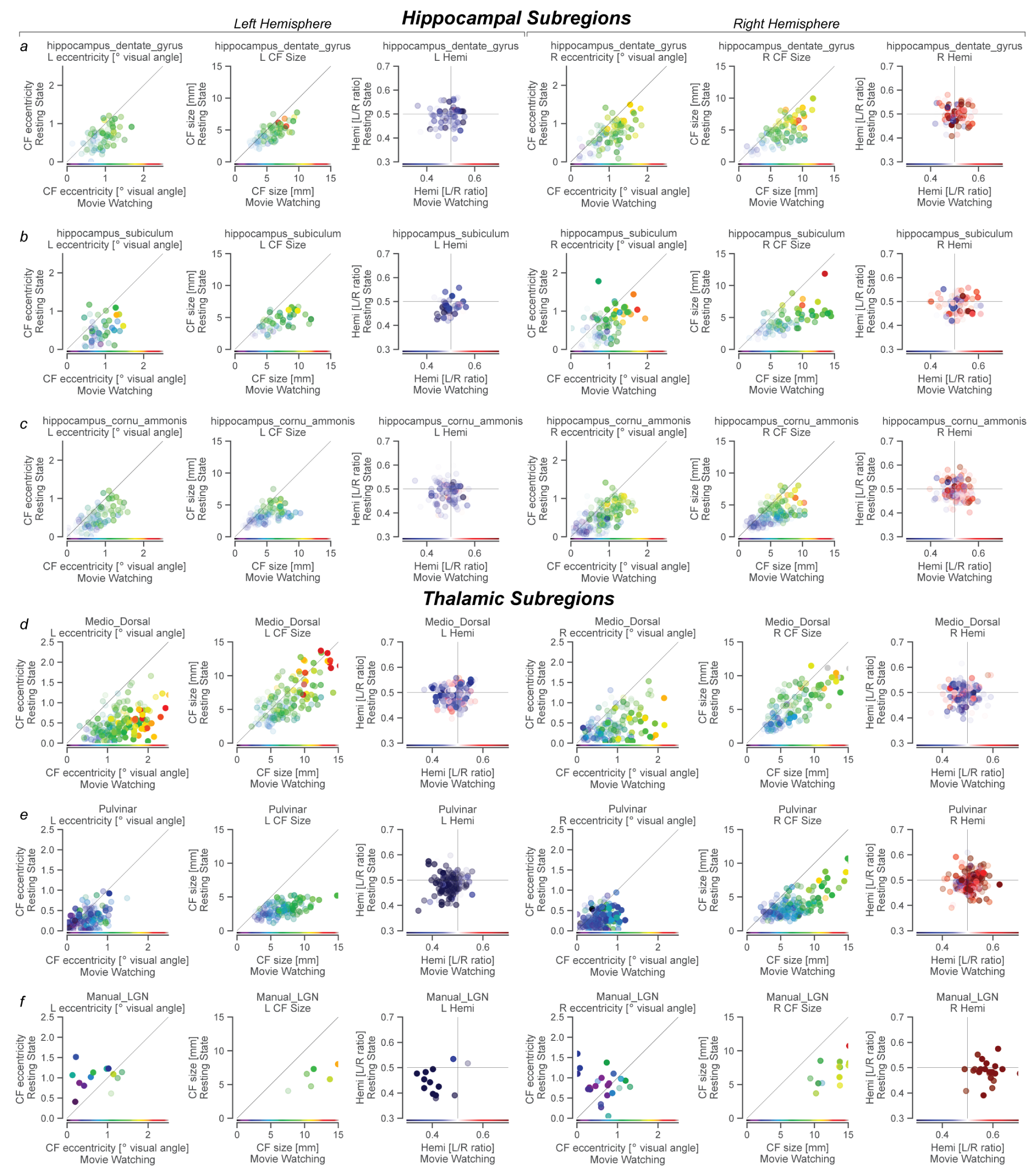
CF parameters in selected subfields of subcortical (Hippocampus and Thalamus) structures, per hemisphere. ROIs taken from Jülich histological and Najdenovska atlases of hippocampus and thalamus, respectively, with LGN delineated manually.

## Notes

### Competing Interest Statement

The authors have declared no competing interest.

